# Hybrid Living Capsules Autonomously Produced by Engineered Bacteria

**DOI:** 10.1101/2020.11.23.394965

**Authors:** Daniel P. Birnbaum, Avinash Manjula-Basavanna, Anton Kan, Neel S. Joshi

## Abstract

Bacterial cellulose (BC) has excellent material properties and can be produced cheaply and sustainably through simple bacterial culture, but BC-producing bacteria lack the extensive genetic toolkits of model organisms such as *Escherichia coli*. Here, we describe a simple approach for producing highly programmable BC materials through incorporation of engineered *E. coli*. The acetic acid bacterium *Gluconacetobacter hansenii* was co-cultured with engineered *E. coli* in droplets of glucose-rich media to produce robust cellulose capsules, which were then colonized by the *E. coli* upon transfer to selective lysogeny broth media. We show that the encapsulated *E. coli* can produce engineered protein nanofibers within the cellulose matrix, yielding hybrid capsules capable of sequestering specific biomolecules from the environment and enzymatic catalysis. Furthermore, we produced capsules capable of altering their own bulk physical properties through enzyme-induced biomineralization. This novel system, based on autonomous biological fabrication, significantly expands the functionality of BC-based living materials.

## Introduction

Living systems have evolved the ability to produce materials—from bacterial biofilms to skeletal tissue—that can respond to environmental cues, self-regenerate, and dynamically alter their physical properties. There is great potential for the development of engineered living materials (ELMs) with these same desirable attributes (*1–3*).

Organisms such as bacteria and yeasts can be genetically programmed to execute sensing, computing, memory, and response functions (*4*). Many approaches have been explored for producing ELMs by incorporating these living, programmable units into human-made materials (*5–14*). ELMs have also been fabricated using natural polymers harvested from microbes (*15–21*). But unlike the living materials found in nature, these ELMs require specialized labor and instrumentation to assemble the biomaterial into the desired architecture. To more closely replicate the living materials found in nature, ELMs should be fabricated via self-organization, with the cells simultaneously producing the material and imbuing it with genetically programmable functionality.

One notable example of bulk material production from living cells is bacterial cellulose (BC). When grown in sugar-rich medium, various species of Gram-negative acetic acid bacteria can produce extracellular cellulose with possible yields of >10 grams per liter after a few days of growth (*22*). This is often in the form of a thick, floating mat—known as a pellicle— that forms atop a static liquid culture, growing at the air-water interface. This pellicle consists of a dense network of cellulose fibrils, each ~50 nm wide and up to 9 μm in length, in which the BC-producing bacteria are embedded (*23*).

BC is both highly pure and highly crystalline, which affords it excellent mechanical properties, including a Young’s modulus of 114 GPa for an individual fibril (*24*). It is biocompatible and has a high capacity for water retention, which, along with its purity, has made it an attractive material for wound and surgical dressings. Furthermore, it is biodegradable and can be mass-produced at low cost, with little environmental impact or necessary equipment. These properties have inspired a wide range of applications including wound dressings (*25*), scaffolds for tissue engineering (*26–28*), electronic paper displays (*29*), substrates for enzyme immobilization (*30, 31*), and acoustic diaphragms for speakers (*32*).

Several methods have been developed to better control the shape, and improve the functionality of BC materials. Greca *et al.* exploited the tendency of BC-producing bacteria to synthesize cellulose at the air-water interface to produce highly customizable, self-assembled 3-D structures (*33*). Song *et al.* utilized this same principle to develop an emulsion-based technique for high-throughput production of BC capsules across a wide range of sizes (*34*). Additionally, several groups have developed methods for genetically engineering BC-producing bacteria to produce BC materials with the ability to dynamically respond to their environment. Florea *et al.* genetically engineered a BC-producing strain so that cellulose production and red fluorescent protein (RFP) production could be controlled by the addition of a small molecule inducer, which allowed for spatial patterning of the cellulose pellicle (*35*). Walker *et al.* genetically engineered “sender” and “receiver” strains of a BC-producing bacteria to achieve boundary detection in a fused pellicle (*36*). Even with encouraging early efforts in this area, the potential functionality of smart ELMs composed of BC is limited by our ability to engineer BC-producing bacteria to secrete recombinant proteins and sense specific external cues.

Yet, by co-culturing BC-producing bacteria with other microorganisms, it is possible to significantly expand the capabilities of BC-based ELMs. Das *et al.* co-cultured BC-producing bacteria with the photosynthetic microalgae *Chlamydomonas reinhardtii* in a symbiotic consortium wherein the oxygen produced by *C. reinhardtii* allowed for increased cellulose production away from the air-water interface (*37*). Recently, Gilbert *et al.* developed a co-culture system in which the model eukaryote *Saccharomyces cerevisiae* is stably maintained among BC-producing bacteria during cellulose production. The yeast provides an engineerable host cell within the growing material that can be rationally programmed at the genetic level for dedicated tasks (*38*). This colocalization of different cell types with specialized functions is common in naturally occurring living materials, and could offer their engineered counterparts a comparable level of versatility and robustness.

In this work, we have developed a novel platform in which a BC-producing strain of bacteria (*Gluconacetobacter hansenii*) is co-cultured with various engineered strains of *E. coli*. By incorporating the engineered *E. coli* within the BC material, we significantly expand the functionality of BC-based ELMs by exploiting some of the many genetic tools developed for *E. coli*. Our approach utilizes the BC-producing *G. hansenii* as a primary “material factory,” while the engineered *E. coli* provides genetically encoded functionality. Through simple steps of incubation and media exchange, we produce BC capsules containing dense colonies of engineered *E. coli* that can respond to environmental cues. We show that the encapsulated *E. coli* can provide functionality by expressing engineered curli nanofibers, yielding hybrid capsules capable of biomolecule sequestration and enzymatic catalysis. Additionally, we demonstrate the ability of the engineered capsules to modulate their own bulk physical properties through urease-induced biomineralization.

## Results

### Biological fabrication of BC capsules containing engineered *E. coli*

We chose to produce the BC material as a hollow spherical capsule rather than the more conventional pellicle format in the hopes of creating a material with topologically distinct regions, with the *G. hansenii* primarily occupying the “outer” shell composed of a dense cellulose matrix, and *E. coli* occupying the “inner” core. To produce the BC capsules, we inoculated Hestrin-Schramm (HS) media with *G. hansenii* and incubated droplets of the culture on a bed of superhydrophobic polytetrafluoroethylene (PTFE) powder, as previously described (*33*). After several days of incubation these droplets became robust BC capsules due to cellulose production by *G. hansenii* at the air-water interface. To produce capsules containing engineered *E. coli*, we inoculated the HS media with both *G. hansenii* and *E. coli*. Then, following the incubation in HS, the newly-formed capsules were transferred to lysogeny broth (LB) media (containing antibiotic to select for engineered *E. coli*) and incubated under shaking conditions to proliferate *E. coli* inside the capsules (Fig. 1A). We lowered the HS media pH from 5.9 to 5 using citric acid because a preliminary analysis showed that *E. coli* survived, but did not proliferate, and that cellulose production from *G. hansenii* was strong at this pH level (fig. S1).

**Fig. 1.**
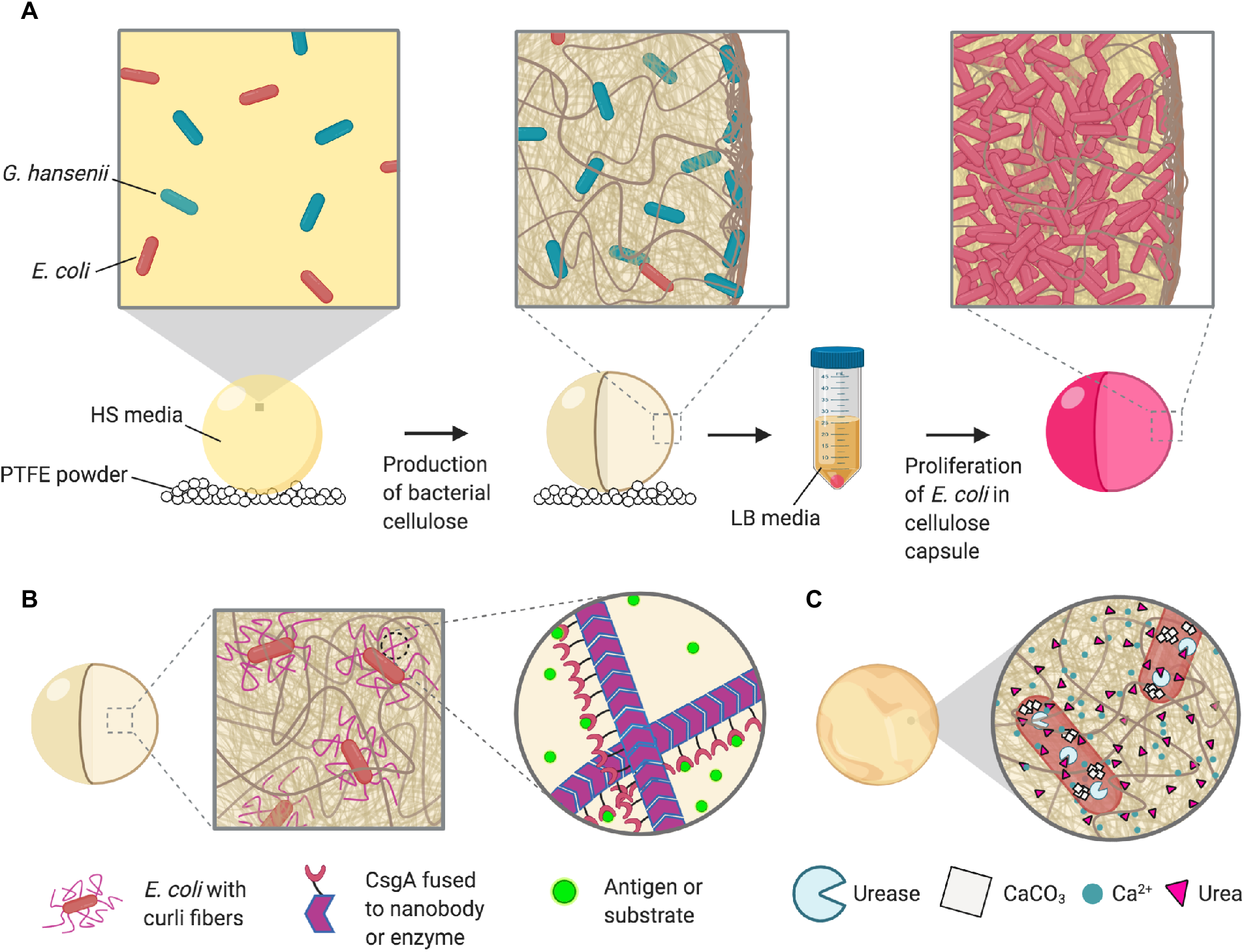
Production of programmable hybrid living materials from bacterial co-culture. **(a)** Schematic showing co-culture in HS media leading to production of bacterial cellulose, followed by transfer to LB media for selective proliferation of *E. coli* inside the BC capsule. **(b)** Schematic showing functionalization of the capsule via production of engineered curli fibers by the encapsulated *E. coli*. **(c)** Schematic showing biomineralization of the capsule through production of urease by the encapsulated *E. coli* leading to CaCO_3_ crystal growth when in the presence of urea and Ca^2+^.

To clearly demonstrate the programmable function of the encapsulated *E. coli*, we used the *E. coli* strain BL21 containing the plasmid pBbA8k-RFP encoding arabinose-inducible expression of mRFP1 (*39*). After the overnight incubation in LB, the capsules inoculated with *E. coli* had a more opaque appearance compared to the capsules without *E. coli* (Fig. 2A) due to the growth of *E. coli* within the capsules. When incubated in LB with the addition of 0.1% arabinose, the capsules exhibited a bright red appearance due to the production of RFP in the engineered *E. coli*. This demonstrated that the plasmid was functional in the encapsulated *E. coli*, and allowed us to clearly visualize the growth of *E. coli* colonies inside the capsules.

**Fig. 2.**
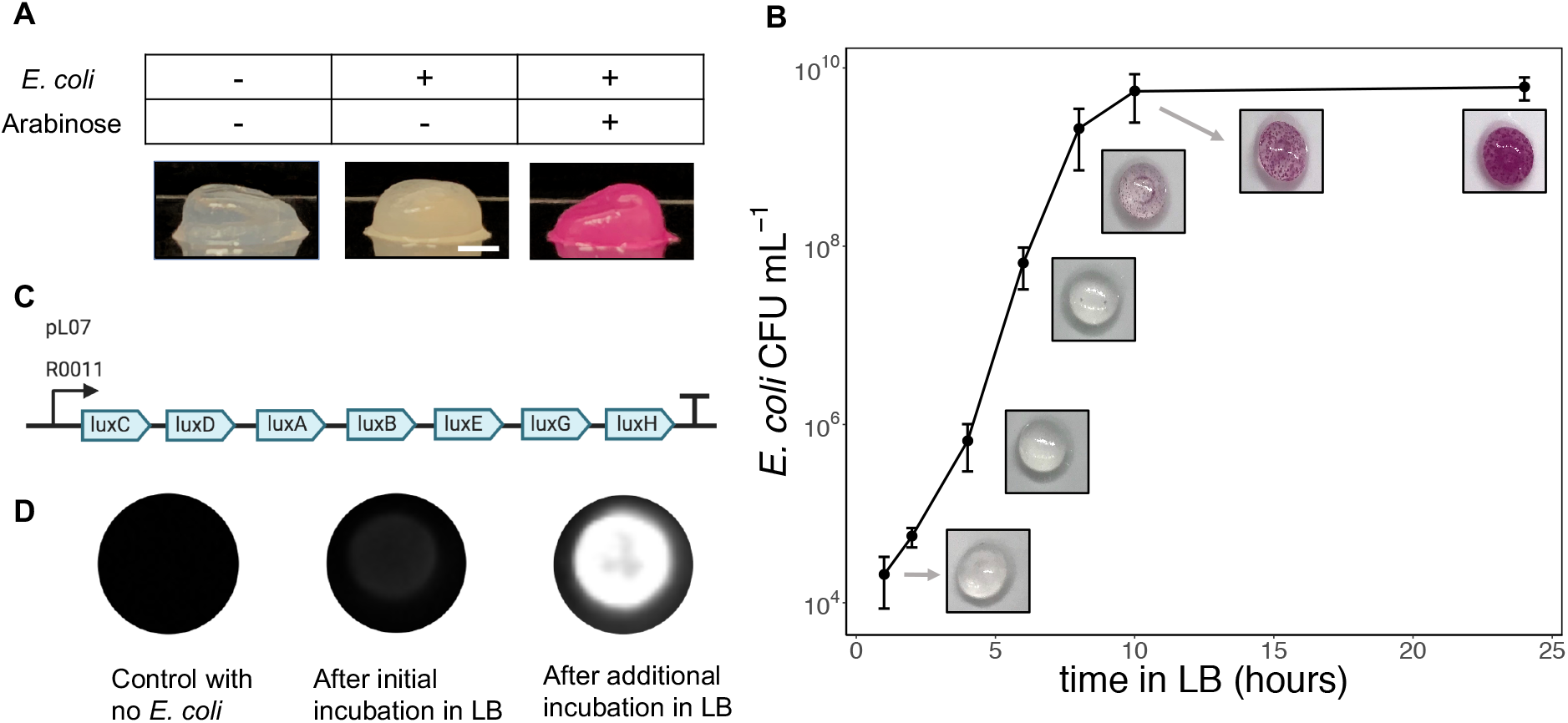
BC capsules containing a high concentration of programmable living *E. coli*. **(a)** Images of BC capsules with and without engineered *E. coli*. The addition of arabinose during the LB incubation step induced RFP expression in the *E. coli*. Scale bar = 2 mm. **(b)** Growth curve of encapsulated *E. coli* generated by degrading capsules after various incubation times in LB and plating on selective LB-agar to quantify CFU. Data on *E. coli* cell density within the capsules is accompanied by images of the BC capsules containing RFP-expressing *E. coli*. **(c)** Plasmid pLO7 encodes constitutive expression of the *luxCDABEGH* operon. **(d)** Capsules containing *E. coli* transformed with pLO7 were stored in PBS overnight following the initial incubation in LB. Some capsules were then incubated for an additional two hours in fresh LB.

We used this RFP-expressing strain to track the progression of *E. coli* growth inside the capsules during the incubation in LB (Fig. 2B). After 6 hours, small *E. coli* colonies became very faintly visible. After 8 hours, many more colonies were visible, with some appearing to be located near the outer wall of the capsule, and others appearing to be located in the core of the capsule. The core colonies often took the form of a ring, possibly due to the rotational motion of the shaking incubator. After 10 and 24 hours, the colonies became larger and more numerous, eventually covering most of the capsule surface. We also measured the growth rate of the *E. coli* during the incubation in LB by degrading the capsules through mechanical homogenization and cellulase treatment, and plating the degradant on selective LB-agar to quantify *E. coli* colony-forming units (CFU) inside the capsules (Fig. 2B). The resulting growth curve showed that the encapsulated *E. coli* grew to a high density, close to 10^10^ CFU mL^−1^, after the overnight incubation in LB.

We wanted to test whether the encapsulated *E. coli* were capable of further activity when the media was replenished. To demonstrate that the encapsulated *E. coli* were metabolically active, we used the plasmid pLO7, which contained a constitutively expressed *luxCDABEGH* operon from the marine bacterium *V. harveyii* (Fig. 2C). The luminescent reaction, unlike a fluorescent protein, requires constant cellular energy, so can only be maintained while the *E. coli* are in a metabolically active state (*40*). After the initial overnight incubation in LB, capsules containing *E. coli* with pLO7 showed a weak luminescence signal. The capsules were then stored in phosphate buffered saline (PBS) overnight before incubation in fresh LB for an additional two hours. During this second incubation in LB, the capsules became strongly luminescent, indicating that the encapsulated *E. coli* were viable and capable of quickly entering a metabolically active state (Fig. 2D).

### Co-culture dynamics during capsule production

The bright red appearance of the RFP-expressing *E. coli* colonies allowed us to visually observe the patterns of *E. coli* growth inside the capsules. We found that both the extent of *E. coli* growth during the incubation in LB and the permeability of the capsules varied significantly depending on at least two factors: the initial inoculation of *G. hansenii* and the duration of incubation in HS media before transfer to LB.

To assess the influence of these factors on *E. coli* growth, we synthesized capsules where the volume of week-old *G. hansenii* starter culture used to inoculate the HS co-culture was varied between 5% and 20% of the total co-culture volume, and the incubation time in HS was varied between 2 and 6 days. Given a 5% inoculation of *G. hansenii*, the *E. coli* survived well during the incubation in HS, as evidenced by the growth of many *E. coli* colonies during the incubation in LB (Fig. 3A). When the inoculation volume was increased to 10%, the survivability of the *E. coli* was reduced, with capsules showing no evidence of *E. coli* growth after 6 days of incubation in HS. When the inoculation volume was increased further to 20%, the decrease in *E. coli* survival was even more significant, with no *E. coli* colonies apparent in capsules incubated for more than 3 days in HS. For all three inoculations, when the incubation in HS was only 2 days long, the capsules were noticeably less robust, and their liquid contents quickly leaked out when placed on a glass slide.

**Fig. 3.**
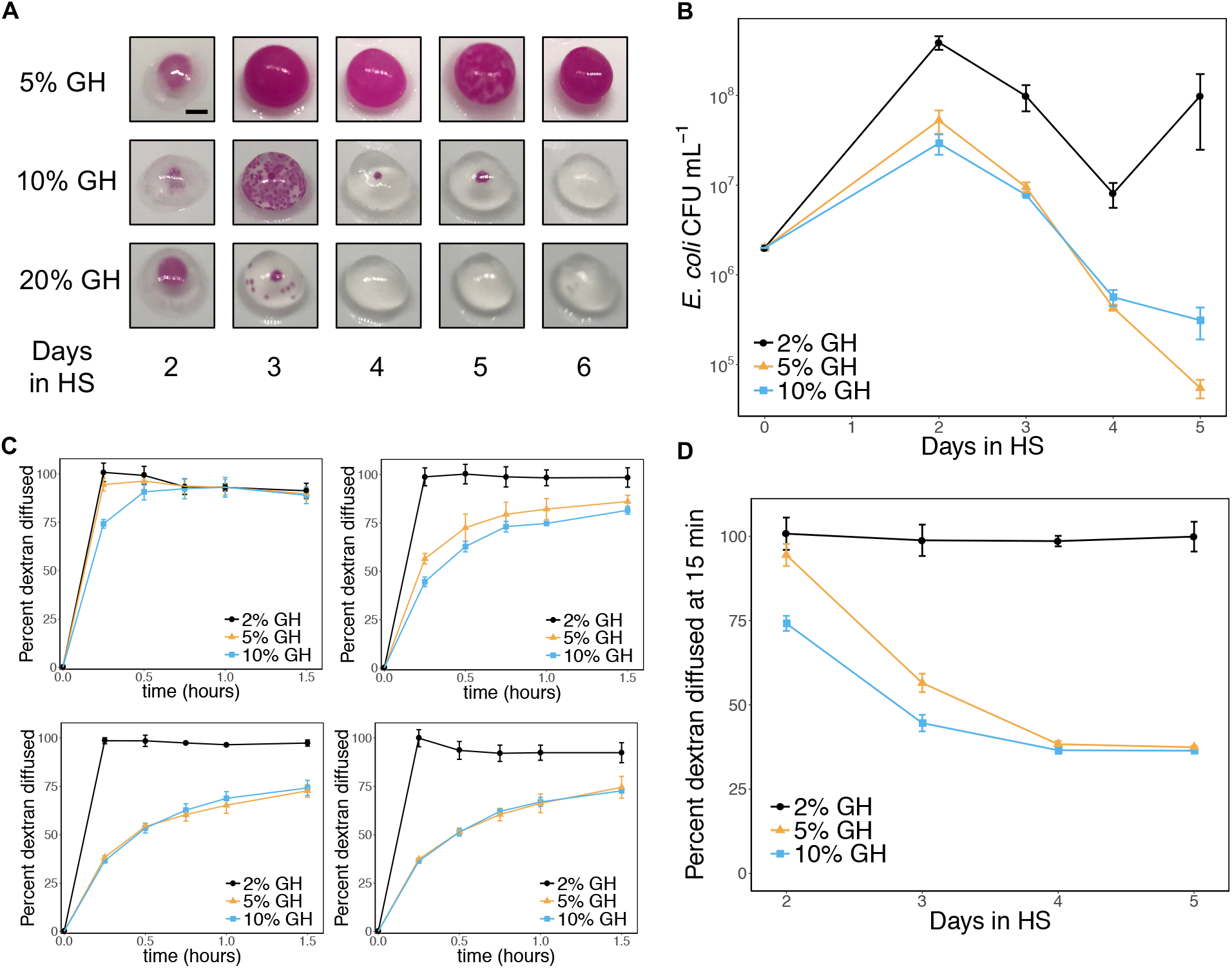
Effects of co-culture parameters on *E. coli* survival and capsule permeability. GH refers to the relative volume of *G. hansenii* starter culture added to the HS co-culture. **(a)** Images of BC capsules containing RFP-expressing *E. coli*. Scale bar = 2 mm. **(b)** Survival of *E. coli* in capsules during the co-culture in HS. After the incubation in HS, the capsules were degraded and plated on selective LB-agar to quantify *E. coli* CFU. **(c)** Diffusion of 500 kDa, FITC-conjugated dextran out of the capsules. HS co-cultures were additionally inoculated with dextran, and capsules were transferred to PBS after the HS incubation period. Permeability was assessed by measuring the FITC fluorescence of the surrounding solution over time. Top left: 2 days in HS. Top right: 3 days in HS. Bottom left: 4 days in HS. Bottom right: 5 days in HS. **(d)** Rate of dextran diffusion. The rate of dextran diffusion out of the capsules is estimated using the fluorescence of the supernatant after the capsules were incubated in PBS for 15 minutes.

To better track *E. coli* survival during the co-culture in HS, we degraded the capsules prior to incubation in LB and plated the degradant on selective LB-agar to quantify *E. coli* CFU. For this experiment, the inoculation of week-old *G. hansenii* starter culture was varied between 2% and 10% of the total co-culture volume. Given a 5% or 10% inoculation of *G. hansenii*, the *E. coli* CFU grew by about an order of magnitude after 2 days in HS, then consistently decreased thereafter (Fig. 3B). When the inoculation volume was lowered to 2%, *E. coli* CFU increased by about two orders of magnitude after 2 days in HS, then decreased for the next two days, before increasing again after 4 days.

Both experiments indicate that increasing the *G. hansenii* inoculation tends to decrease the number of surviving *E. coli* available to colonize the capsule upon incubation in LB. This suggests that the two microorganisms compete for resources during the co-culture in HS media. For most of the *G. hansenii* inoculations tested, the number of surviving *E. coli* also tended to decrease with longer incubation times in HS. However, when the *G. hansenii* inoculation was lowered to 2% in the second experiment, the *E. coli* concentration rebounded on the fifth day in HS, reflecting the importance of the relative initial concentrations of the two bacteria on the competition dynamics.

Next, to evaluate the effects of the co-culture dynamics on the permeability of the capsules, we measured the rate at which high molecular weight dextran diffused out of the capsules. For this experiment, the HS media was inoculated with 0.5 mg/mL fluorescein isothiocyanate (FITC)-conjugated-, 500 kDa dextran, in addition to *G. hansenii* and *E. coli*. The inoculation of *G. hansenii* was again varied between 2% and 10% of the total co-culture volume. After the incubation in HS, the capsules were transferred to 5 mL PBS, and the rate at which dextran diffused out was measured by monitoring the fluorescence of the surrounding solution (Fig. 3C). For capsules inoculated with 5% or 10% *G. hansenii*, the rate of dextran diffusion decreased significantly as HS incubation time was increased from 2 to 5 days (Fig. 3D), likely because of a denser cellulose matrix. In contrast, when the *G. hansenii* inoculation was 2%, the rate at which dextran escaped from the capsules did not decrease even after 5 days of incubation in HS. This could indicate that cellulose production in these capsules was hindered by the relatively low inoculation of *G. hansenii*. In all cases, the dextran diffused out of the capsules within several hours, demonstrating their permeability to a wide size range of biological molecules.

### Internal structure of BC capsules containing engineered *E. coli* colonies

To better understand the interactions between the cellulose matrix produced by *G. hansenii* and the engineered *E. coli*, we imaged thin sections of the capsules using fluorescence microscopy (Fig. 4 A, B, and C). The cellulose matrix was stained with Calcofluor White (CFW), a blue fluorescent dye which binds cellulose, while the *E. coli* fluoresced due to RFP expression. To maintain the 3-D structure of the mostly hollow samples, including the shape and distribution of the *E. coli* colonies, we equilibrated the capsules in gelatin then fixed them in paraformaldehyde (PFA) prior to cryosectioning.

**Fig. 4.**
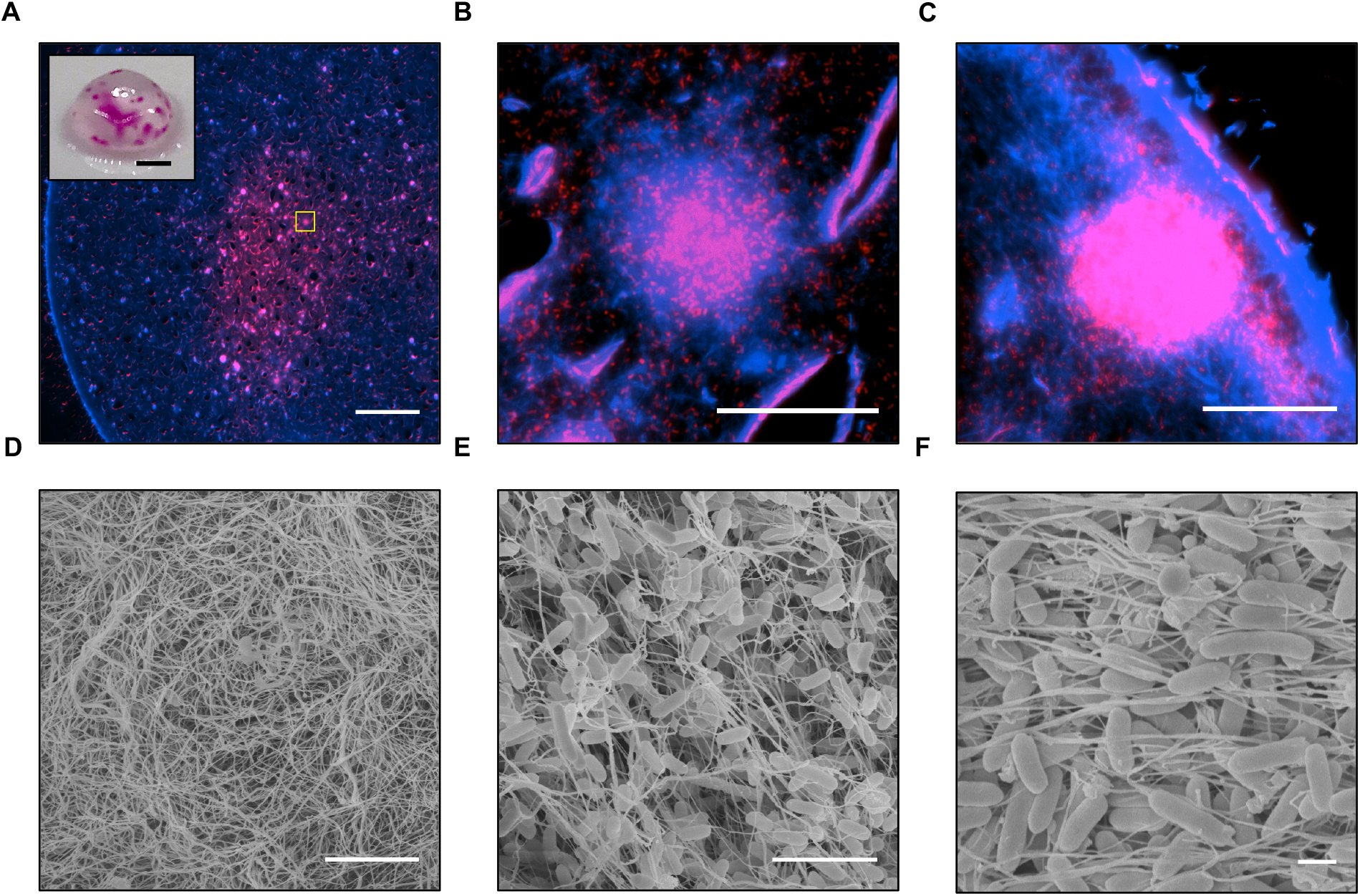
Internal structure of BC capsules containing engineered *E. coli*. Panels **(a) – (c)** show 8 μm cross-sections of a capsule pre-stained with cellulose stain Calcofluor White (CFW). CFW fluorescence is shown in blue and RFP fluorescence is shown in red. **(a)** A large region of *E. coli* growth is visible in the core of the capsule. The outer wall of the capsule is visible as a bright blue band. Scale bar = 500 μm. Inset: a photograph of the capsule taken before staining with CFW; scale bar = 2 mm. **(b)** Zoomed-in view of the region in the yellow box in panel (a). Scale bar = 50 μm **(c)** View of an internal *E. coli* colony located against the outer wall of the capsule. Scale bar = 50 μm. Panels (**d) – (f)** show FESEM images of capsules. **(d)** View of the outer wall of the capsule, showing a dense network of cellulose fibers. Scale bar = 5 μm. **(e)** View of encapsulated *E. coli* enmeshed within the internal cellulose matrix. Scale bar = 5 μm. **(f)** Another view of internal, encapsulated *E. coli* densely packed together. Scale bar = 1 μm.

Visual inspection of the capsule prior to sectioning showed both a large internal colony and several smaller colonies along the outer wall (Fig. 4A, inset). Imaging of the sections revealed that the large internal colony was not homogenous in cell density, but was comprised of many smaller, highly concentrated colonies connected by more diffuse regions of *E. coli* (Fig. 4A). At higher magnification, we found that the highly concentrated pockets of *E. coli* corresponded to regions of high cellulose fiber density (Fig. 4B).

Imaging of the sections also revealed an *E. coli* colony located along the outer wall of the capsule (Fig. 4C). This likely corresponds to one of the colonies forming dots on the surface of the capsule. The dense outer wall of the capsule, visible as a bright blue band roughly 20 μm thick, seemed to mostly contain this concentrated pocket of *E. coli*. The localization of both this colony and the more internal colonies in regions dense with cellulose fibers suggests that the *E. coli* may use the cellulose matrix as a solid substrate for growth within the capsule.

Additionally, we noted that the preparation and sectioning of the capsules led to the formation of apparent “rips” in the imaged sections. These are likely artifacts of the sample preparation and cryosectioning procedures, possibly due to non-uniform embedding in the gelatin or tearing of the tissue during sectioning. We noted that the gelatin-embedding procedure did not affect the visual appearance of the capsules, preserving the macroscopic features of the cellulose sphere. As such, we felt this technique adequately captured the spatial distribution of the dense bacterial populations within the capsules.

To further elucidate the structure of the capsules, we performed field emission scanning electron microscopy (FESEM) imaging of capsules cut open with a scalpel. Images of the exterior of the capsule showed the dense but porous nature of the outer cellulose wall (Fig. 4D). Images of the capsule interior showed *E. coli* growing along the cellulose matrix (Fig. 4E) and in tightly-packed conformations (Fig. 4F).

### Functional hybrid capsules containing engineered curli nanofibers

To demonstrate the potential functionality of the capsules, we produced hybrid capsules containing engineered curli nanofibers displaying various functional protein domains. Curli are insoluble and robust functional protein fibers anchored to the surface of *E. coli* cells, whose versatility as a platform for ELMs has been explored in many applications, including adhesion to abiotic surfaces (*16*), *in vivo* display of therapeutic domains (*41*), and sequestration of pathogens from drinking water (*42*), among others (*43*). We hypothesized that we could add a range of functionality to the capsules by expressing curli-tethered proteins, which would remain fixed within the BC matrix while interacting with the external solution.

First, we attempted to use the hybrid capsules to sequester specific proteins from surrounding solution. We used *E. coli* engineered to express curli fibers in which the major curli subunit CsgA was fused to a single-domain antibody (sometimes known as a “nanobody”) specific for green fluorescent protein (NbGFP) (Fig. 1B) (*45*). Plasmid pBbB8k-csg-NbGFP, which contained an arabinose-inducible synthetic curli operon *csgBACEFG* with CsgA fused to NbGFP via a flexible linker, was transformed into curli knockout strain PQN4 (Fig. 5A). When curli production was induced, the NbGFP-expressing capsules sequestered over 1 μg of GFP per capsule from a purified GFP solution (Fig. 5B). When the capsules contained curli fibers displaying an alternate antibody domain NbStx2, which is specific for Shiga toxin 2 (*46*), GFP was not sequestered, demonstrating the specificity and efficacy of NbGFP.

**Fig. 5.**
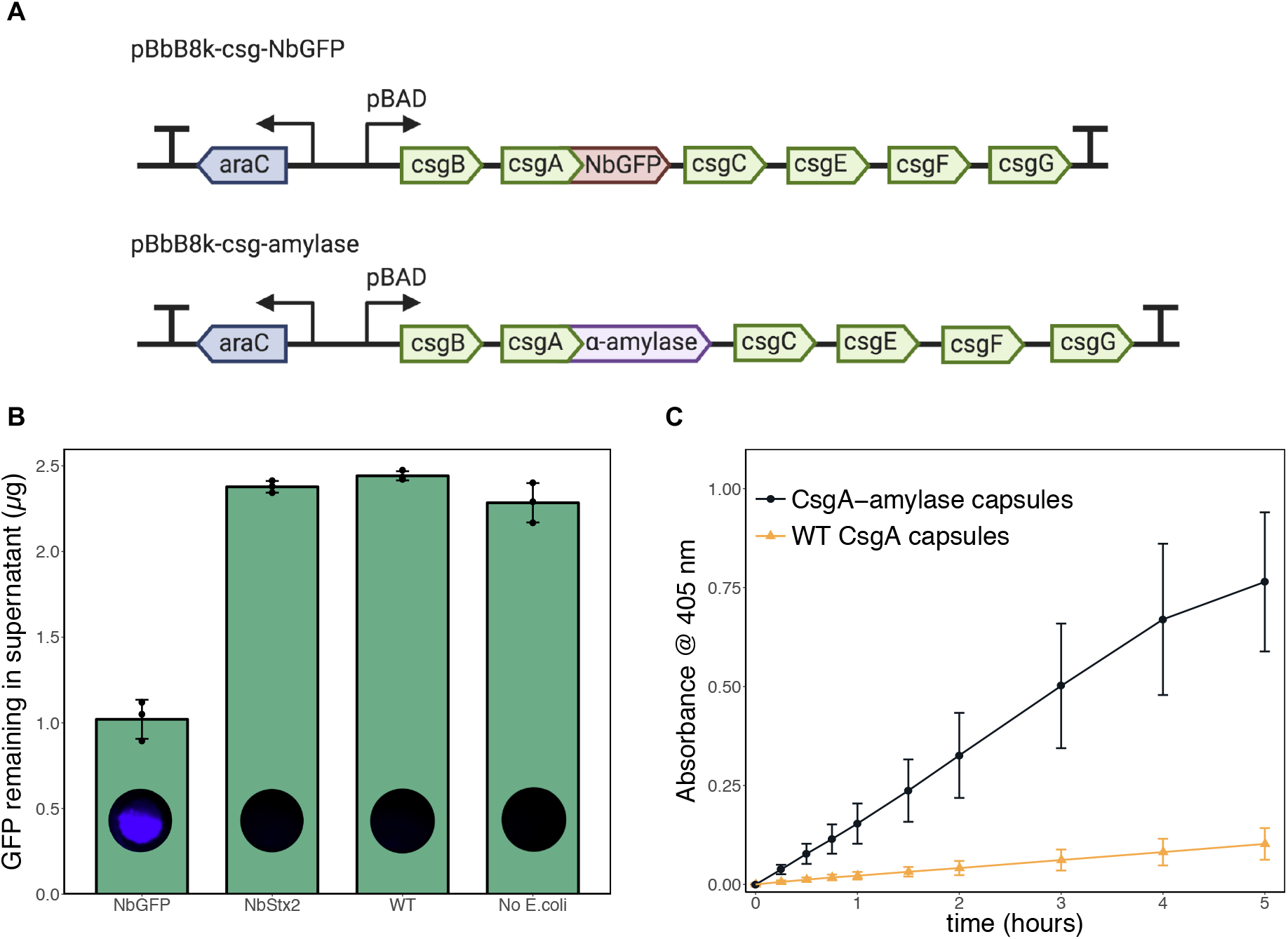
Functional hybrid capsules containing engineered curli nanofibers. **(a)** Plasmids encoding arabinose-inducible expression of engineered curli fibers. **(b)** Bar graph showing green fluorescent protein (GFP) levels after overnight incubation of capsules in 0.5 mL of purified 4.6 μg/mL GFP solution. When the capsules contained curli fibers displaying a nanobody domain specific for GFP, there was a significant reduction in the GFP concentration. After incubation in GFP solution, capsules were washed and imaged for GFP fluorescence. **(c)** Time course of α-amylase activity in capsules, measured by incubation of the capsules with the chromogenic substrate 4-nitrophenyl α-D-maltohexaoside, which produces a yellow solution upon cleavage by α-amylase.

As another demonstration of capsule functionality, we encapsulated *E. coli* cells in which CsgA was fused to a model enzyme—α-amylase from the soil bacterium *Bacillus licheniformis*. α-Amylases catalyze the hydrolysis of starch molecules into smaller oligosaccharides. They have applications in the food, fermentation, textile, paper, detergent and pharmaceutical industries (*47*). α-Amylase activity was assessed by incubating the capsules in the presence of a model substrate, 4-nitrophenyl α-D-maltohexaoside, which produces a yellow product upon cleavage by α-amylases (*48*). We observed significantly more enzyme activity in capsules containing curli fibers with α-amylase compared to capsules containing wild-type CsgA curli fibers (Fig. 5C). After 5 hours of incubation in the presence of the model substrate, a single capsule’s amylase activity was equivalent to roughly 400 mU of purified amylase (fig. S5).

### Programmable biomineralization of capsules

One hallmark of natural living materials is the ability to alter their own bulk physical properties. To demonstrate this ability in our system, we sought to produce capsules capable of biomineralization. To achieve this, we used a commercially available *E. coli* strain, HB101, transformed with the plasmid pBR322-Ure, which contains the urease gene cluster from the soil bacterium *S. pasteurii* (*49*). Urease catalyzes hydrolysis of urea, which leads to the formation of carbonate ions and an increase in pH. In the presence of soluble calcium ions, this leads to the precipitation of CaCO_3_ (Fig. 1C).

When capsules containing HB101 / pBR322-Ure cells were incubated in growth media supplemented with urea and CaCl_2_, levels of soluble calcium in the culture supernatant decreased significantly compared to cultures containing capsules with HB101 cells without a plasmid, presumably due to urease-induced precipitation of CaCO_3_ (Fig. 6A). Incubation in the urea and CaCl_2_-containing media caused a drastic change in the physical appearance of the HB101 / pBR322-Ure capsules, which also exhibited a striking increase in stiffness compared to capsules containing HB101 cells without a plasmid (Fig. 6B). We observed the mineralization process by imaging capsules incubated in the urea and CaCl_2_-containing media for various amounts of time. We saw that the capsules changed in appearance quite rapidly, and took on a yellowish, pebble-like appearance after only 2 hours of incubation (Fig. 6C). By comparison, the capsules containing *E. coli* without pBR322-Ure did not change in appearance during such an incubation.

**Fig. 6.**
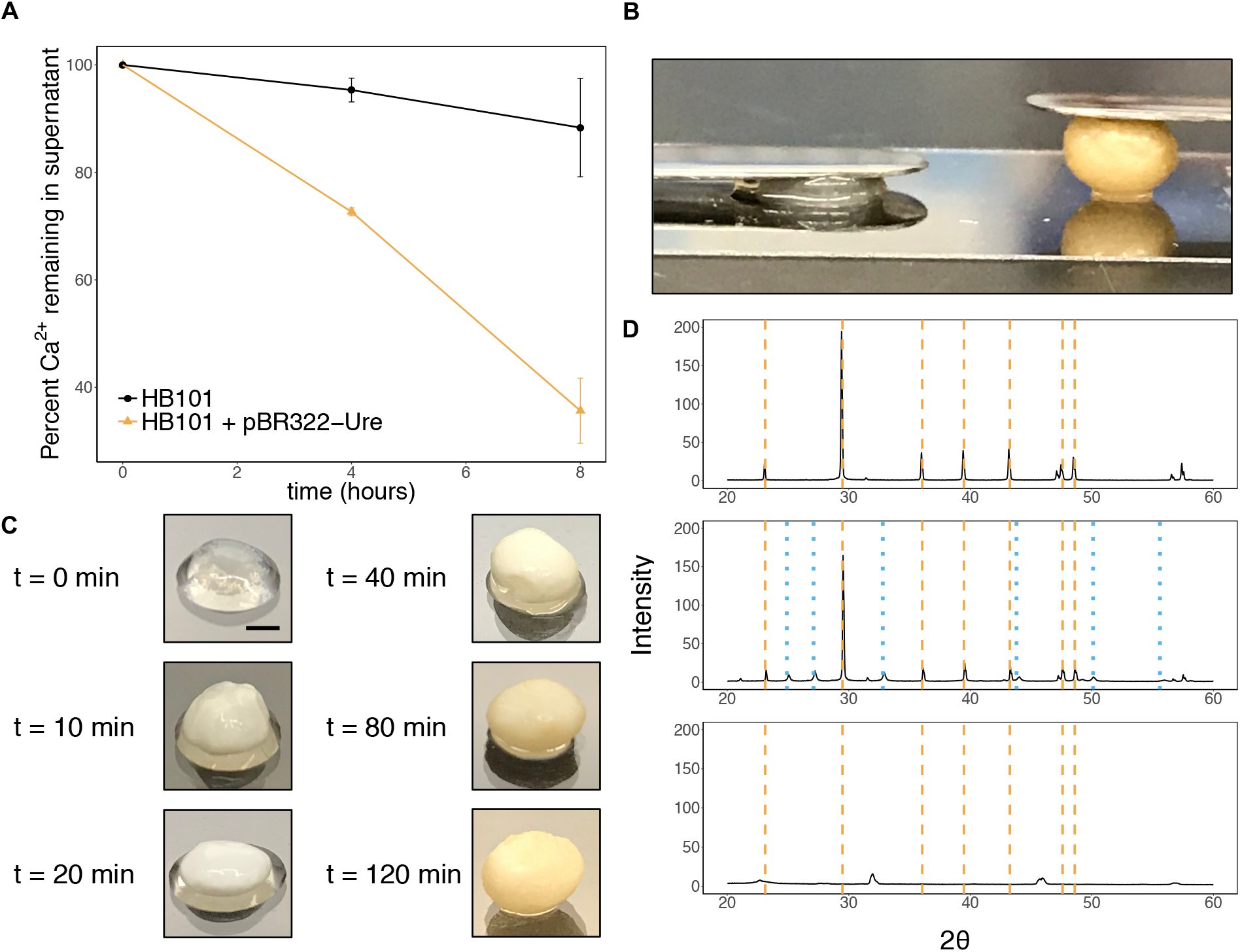
Biomineralization of capsules via urease expression in *E. coli*. **(a)** Soluble calcium levels in the culture supernatant during incubation of the capsules in urea and CaCl_2_-containing media, normalized by initial calcium levels. Calcium levels decreased significantly faster when the capsules contained the HB101/pBR322-Ure cells. **(b)** Image of capsules after an 8-hour incubation in urea and CaCl_2_-containing media, showing an increase in stiffness for the mineralized capsule. Left capsule contained HB101 cells with no plasmid, and right capsule contained HB101 cells with plasmid pBR322-Ure encoding the urease gene cluster from *S. pasteurii*. Each capsule has a 7 g stainless steel spatula resting on it. **(c)** Images of HB101 / pBR322-Ure capsules incubated in urea and CaCl_2_-containing media for various amounts of time. Scale bar = 2 mm. **(d)** XRD patterns. After the incubation in urea and CaCl_2_-containing media, capsules were dried at room temperature and ground into a fine powder using a mortar and pestle prior to XRD analysis. Top: precipitate from a culture of non-encapsulated HB101 / pBR322-Ure cells. Middle: Capsules containing HB101 / pBR322-Ure capsules. Bottom: Capsules containing HB101 with no plasmid. Dashed orange lines indicate peaks for calcite, and dotted blue lines indicate peaks for vaterite (*50–51*).

To confirm the presence of CaCO_3_ mineral forms in the yellowed capsules, we performed X-ray powder diffraction analysis (XRD). After the incubation in the urea and CaCl_2_-containing media, the capsules were left to dry at room temperature, and then ground into a fine powder using a mortar and pestle. As a control, we also analyzed precipitate from a culture of non-encapsulated HB101 / pBR322-Ure cells. The analysis showed that the precipitate from the non-encapsulated cell culture contained mostly calcite (Fig. 6D). Interestingly, while the mineralized capsules indeed contained calcite, the XRD profile also indicated the presence of vaterite, another CaCO_3_ mineral form. This suggests that the presence of the cellulose matrix significantly impacts the mineralization process.

## Discussion

We have developed an approach involving only simple microbial culture steps to produce robust, programmable materials containing a high concentration of engineered, living cells. Our co-culture system leverages the strengths of two different microbes by enabling BC-producing bacteria to produce a robust cellulose matrix colonized by programmable engineered *E. coli*. Since *E. coli* are known to have a much faster growth rate than *G. hansenii*, we first incubated the co-culture in pH-lowered HS media to suppress *E. coli* growth and encourage cellulose production from *G. hansenii*, then transferred the capsules to LB to proliferate the engineered *E. coli*. Our work significantly extends the capability of BC-based materials by leveraging the extensive genetic engineerability of *E. coli*.

We were initially motivated to incubate the co-culture as spherical droplets in the hopes that the dense cellulose matrix would prevent *E. coli* escape into the surrounding environment. Unfortunately, during the incubation in LB, the culture usually became turbid due to *E. coli* cells escaping from within the capsule. Despite this, the spherical shape of the capsules offers several advantages over the more conventional flat pellicle format. The dense outer wall of the capsule provides a protective barrier for the cells and other possible cargo contained within, while still allowing macromolecular diffusion. Relatively large cargo, such as microparticles, could easily be loaded into the capsules by inoculation into the initial co-culture, and be stably contained by the capsule wall thereafter. The encapsulation approach also ensures the *E. coli* are stably incorporated in the BC material. Moreover, the capsules are quite robust and can be easily handled and transported without breaking.

The extensive genetic engineerability of *E. coli* facilitates the fabrication of living materials with many potential functions. In this work, we showed that robust BC capsules can be genetically programmed to respond to stimuli from the environment. We also used curli fiber-based display to produce hybrid capsules containing engineered functional domains. Using this approach, we produced capsules containing single-domain antibodies that could specifically sequester GFP from solution. Since BC is a robust, biocompatible material, these capsules could be used for the sequestration of specific biomolecules from a variety of complex biological samples. We also used curli fiber display to integrate α-amylase, an industrially relevant enzyme used in ethanol production and many other applications, into the capsules. During the growth in LB, the CsgA-enzyme fusion protein is secreted from *E. coli* and immobilized in the capsules through incorporation in cell-anchored curli fibers, circumventing the need for costly protein purification and enzyme immobilization steps. Our approach is also highly adaptable. We demonstrate encapsulation of several different *E. coli* strains expressing recombinant proteins from a range of different organisms. Furthermore, since the capsules can respond to signals from their environment, they can be programmed for various functions and adapted to specific situations.

As highlighted above, we have demonstrated that the capsules can support immobilized enzymes for catalysis or antibody domains for sequestration. However, this platform would likely require significant development and optimization depending on the intended application. One limitation is the exposure of the genetically engineered *E. coli* to the environment, which precludes this technology from implementation in open environmental settings. Various biocontainment strategies have been developed to circumvent this issue, including genetic safeguards, such as engineered auxotrophy (*52*), and physical containment (*53*). Alternatively, one could sterilize the capsules after production of the relevant functional protein. Another potential limitation of the system is the yield of secreted protein. Though the curli system offers certain advantages, such as multivalent display of the functional domain, over-expression of the curli operon causes a significant decrease in growth, limiting the amount of protein that can be secreted. However, our platform benefits from the many genetic tools for *E. coli*, so that other approaches such as cell surface protein display (*54*), or the secretion of proteins tagged with cellulose-binding domains (*55*), could be used to functionalize the BC capsules.

The three-dimensional BC capsules described here are produced spontaneously, under mild conditions, and through simple biological processes that largely occur autonomously. No exogenous polymers were added to the system to provide a template for growth or to reinforce the mechanical properties of the final product. The straightforward and economical fabrication of the capsules could easily be implemented in many facilities. Given the extensive set of well-characterized genetic parts designed specifically for *E. coli* and the simplicity of our approach, we believe this work represents a significant step forward in the design and fabrication of both functional BC materials and ELMs that mimic the autonomous structure building programs employed by natural living systems.

## Materials and Methods

### Strains and plasmids used in this study

Strains used in this study are listed in Supplementary Table 1. Plasmids used in this study are listed in Supplementary Table 2. All plasmids from this study were constructed using standard cloning techniques (polymerase chain reaction and one-step isothermal Gibson assembly). Oligonucleotides were obtained from IDT. All plasmids were transformed into Mach1 (Invitrogen C862003) for amplification. Constructs were verified by Sanger sequencing (Genewiz).

### Production of *G. hansenii* starter culture

*G. hansenii* colonies were obtained by streaking the strain ATCC 53582 onto an HS-agar plate. The streaked plate was incubated at 25 °C until rough-edged colonies formed. One of these colonies was used to inoculate 10 mL of liquid HS media. HS media is glucose-rich and standard for the production of BC with bacteria. HS media was prepared with 20 g/L glucose, 5 g/L yeast extract, 5 g/L peptone, 2.7 g/L Na_2_HPO_4_, 1.5 g/L citric acid, and 15 g/L agar where required. The culture was incubated for 7 days at 25 °C, forming a thick cellulose pellicle. To produce capsules, this starter culture was vortexed for 1 minute to shake cells loose from the pellicle. The liquid portion of the vortexed culture was then used to inoculate fresh HS media for capsule production. This procedure generally yielded starter cultures with concentration 2 – 4 million CFU/mL, prior to dilution into the final co-culture.

### Evaluation of media pH on *E. coli* survival and BC production

Before defining a standard protocol for producing BC capsules with encapsulated *E. coli*, we tested how the pH of the HS media would affect *E. coli* survival and BC production. *E. coli* BL21 colonies were inoculated in LB and grown overnight under shaking conditions to produce starter cultures. The starter cultures were diluted to OD_600_ = 0.2 and then diluted 1/100 into 10 mL HS media in a Falcon tube. *G. hansenii* starter cultures were prepared as described above and diluted 1/10 into the co-culture. This was repeated using HS media across a range of pH values, with the pH of the HS media adjusted using 100 mM citric acid, pH 1. Unadjusted HS media had a pH of 5.9. The co-culture was then incubated at 25 °C for 5 days. The tube was thoroughly vortexed for 1 minute and the liquid media underlying the pellicles was plated on selective LB-agar to quantify the *E. coli* concentration. The BC pellicles were removed, washed overnight in 0.1 M NaOH, and washed overnight again in DI water, before being allowed to air-dry at room temperature. The dried pellicles were then weighed.

### Standard protocol for production of BC capsules with encapsulated *E. coli*

HS media was adjusted to pH 5 with citric acid before inoculation with *G. hansenii* and *E. coli* starter cultures. The *E. coli* starter cultures were diluted to OD_600_ = 0.2 and then diluted 1/100 into the final co-culture. *G. hansenii* starter cultures were prepared as described above and diluted 1/10 into the final co-culture (unless otherwise specified). The co-culture was drop-casted in 50 μL volumes onto a bed of PTFE powder, forming “liquid marbles”. These droplets were incubated at 25 °C in a humid environment for 3 days (unless otherwise specified) to produce cellulose capsules. A stainless-steel spatula was used to transfer a single capsule to 5 mL LB with antibiotic to selectively grow engineered *E. coli*. The LB culture was incubated overnight at 37 °C with shaking for 18-24 hours. Notably, in most cases, the LB culture would be turbid after the overnight incubation due to *E. coli* escaping from within the capsule.

### Degradation of capsules

The capsules were homogenized in 0.5 mL sterile PBS at 50 Hz for 10 minutes using a 5 mm stainless-steel ball bearing. Then, 30 μL of aqueous cellulase solution (Sigma-Aldrich #C2730) was added to the mixture at a final concentration of 3% v/v cellulase, corresponding to a concentration of ≥ 23.1 units/mL. The mixture was then incubated at 37 °C for 2 hours. At 30 minute intervals during the 2-hour incubation period, the suspension was homogenized at 50 Hz for 2 minutes to accelerate degradation, including a final round of homogenization at the end of the 2-hour period. The PBS containing the degraded capsule was then plated on selective LB-agar to quantify *E. coli* concentration.

### Measuring diffusion rate of dextran out of capsules

To produce dextran-containing capsules, HS media was inoculated with 0.5 mg/mL FITC-labeled 500 kDa dextran (Sigma-Aldrich #46947) in addition to *G. hansenii* and *E. coli* starter cultures (BL21 / pBbB8k-RFP). The capsules were produced as described above, except after the incubation in HS each capsule was transferred to 5 mL PBS and incubated at 37 °C with agitation. 150uL samples of the supernatant were taken at various time points and FITC fluorescence was measured on a BioTek Synergy NEO plate reader (ex,em = 485,528 nm) to determine the kinetics of dextran diffusion from within the capsule into the surrounding PBS.

### Cross-sectional imaging of capsule

A capsule was produced using the standard protocol described above with the strain BL21 and plasmid pBbB8k-RFP. LB was supplemented with kanamycin and 0.1% arabinose to induce mRFP1 expression. Following growth in LB, the capsule was washed three times in 1 mL PBS to remove loosely attached cells. The capsule was then immersed in Calcofluor White Stain (Sigma-Aldrich #18909) for 2 hours to stain cellulose. The capsule was then embedded in gelatin following a previously established protocol (*44*). This was done to maintain the 3-D structure of the capsule throughout the optimal-cutting temperature (OCT) compound embedding procedure. First, the stained capsule was incubated in 12.5% sterile gelatin (Sigma-Aldrich #G2500) at 37 °C with agitation for 24 hours. The capsule was then transferred to 25% gelatin and incubated for another 24 hours at 37 °C with agitation. The capsule was then incubated at 4 °C for 15 minutes to solidify the gelatin. The solidified capsule was then transferred to 25% gelatin and incubated for 24 hours at 37 °C with agitation. Finally, the capsule was fixed by immersion in 4% PFA for 24 hours at room temperature. The fixed capsule was stored in PBS at 4 °C. 8 μm sections were obtained via OCT embedding and cryosectioning. The sections were imaged using an EVOS fluorescent microscope with a DAPI light cube for CFW (357/44 nm excitation; 447/60 nm emission), and a RFP light cube for mRFP1 (531/40 nm excitation; 593/40 nm emission).

### GFP sequestration assay

Capsules were produced using the standard protocol described above using the strain PQN4 and plasmids pBbB8k-csg-NbGFP, pBbB8k-csg-WT, or pBbB8k-csg-NbStx2. LB was supplemented with kanamycin and 0.001% arabinose to induce expression of the curli operon. After the incubation in LB, the capsules were washed three times in 1mL PBS to remove loosely-attached cells. The capsules were then added to 0.5 mL PBS containing 4.6 μg/mL purified GFP and incubated for 24 hours at 37 °C with agitation. Following this incubation, 150 μL samples of the supernatant were taken and GFP fluorescence was measured on a BioTek Synergy NEO plate reader (ex,em = 485,528 nm) to determine the amount of GFP that had been sequestered inside the capsules. The capsules were then thoroughly washed three times in PBS for a total of 3 hours, and imaged for GFP fluorescence using a FluorChem M imager.

### α-Amylase assay

Capsules were produced using the standard protocol described above with the strain PQN4 and plasmids pBbB8k-csg-amylase or pBbB8k-csg-WT. LB was supplemented with kanamycin and 0.001% arabinose to induce expression of the curli operon. After the incubation in LB, the capsules were washed three times in 1 mL PBS to remove loosely-attached cells. The capsules were then transferred to 0.5 mL PBS containing 3 mg/mL of the substrate 4-Nitrophenyl α-D-maltohexaoside (Sigma-Aldrich #73681). The mixture was then incubated at 37 °C with gentle agitation. 150 μL samples of the supernatant were taken with replacement at various time points and absorbance at 405 nm was measured on a BioTek Synergy NEO plate reader to determine the kinetics of α-amylase activity. The same procedure was conducted using various concentrations of purified α-amylase (500-1500 U/mg, Sigma-Aldrich #A4551) to create a calibration curve.

### Biomineralization of capsules

Capsules were produced using the standard protocol described above with the strain *E. coli* HB101 and plasmid pBR322-Ure, or with no plasmid in the control. LB media was supplemented with carbenicillin and 5 μM NiCl_2_. After the incubation in LB, the capsules were stored overnight in PBS at 4 °C. Urea-containing media was prepared with 3 g/L Difco Nutrient Broth, 2.12 g/L NaHCO_3_, 20 g/L NH_4_Cl, and 40 g/L urea. To induce mineralization, capsules were incubated in urea-containing media supplemented with 5 μM NiCl_2_ and 50mM CaCl_2_ at 37 °C with agitation. Soluble calcium levels in culture supernatant were measured using Calcium Colorimetric Assay kit (Sigma-Aldrich #MAK022).

### X-ray powder diffraction of mineralized capsules

After the incubation in urea-containing media, the mineralized capsules were washed three times in MilliQ water for a total of 3 hours, dried at room temperature, and ground into a fine powder using a mortar and pestle. X-ray diffraction was then carried out using a D2 PHASER (Bruker) with a Cu anode operating at 30 kV and 10 mA with a 2θ range of 20° to 60°, step size of 0.02°, and exposure time of 1 second/step.

### Field Emission Scanning Electron Microscope (FESEM) sample preparation

FESEM samples were prepared by fixing the capsules with 2% (m/v) glutaraldehyde and 2% (m/v) paraformaldehyde at room temperature, overnight. The capsules were gently washed with water, and the solvent was gradually exchanged to ethanol with an increasing ethanol 15-minute incubation step gradient (25, 50, 75 and 100% (v/v) ethanol). The capsules were dried in a critical point dryer, placed onto SEM sample holders using silver adhesive (Electron Microscopy Sciences) and sputtered until they were coated in a 10-20 nm layer of Pt/Pd/Au. Images were acquired using a Zeiss Ultra55/Supra55VP FESEM equipped with a field emission gun operating at 5-10 kV.

## Acknowledgments

We thank Suzanne White (Beth Israel Deaconess Medical Center) for performing cryosectioning. We thank Dr. Emily Freed (University of Colorado Boulder) for providing the pBR322-Ure plasmid. We thank Dr. Shao-Liang Zhang (Harvard University) for performing XRD analysis. This work was funded by the National Science Foundation (DMR 2004875).

