## Supplemental Materials for "Hybrid Living Capsules Autonomously Produced by Engineered Bacteria"

### Supplementary Materials

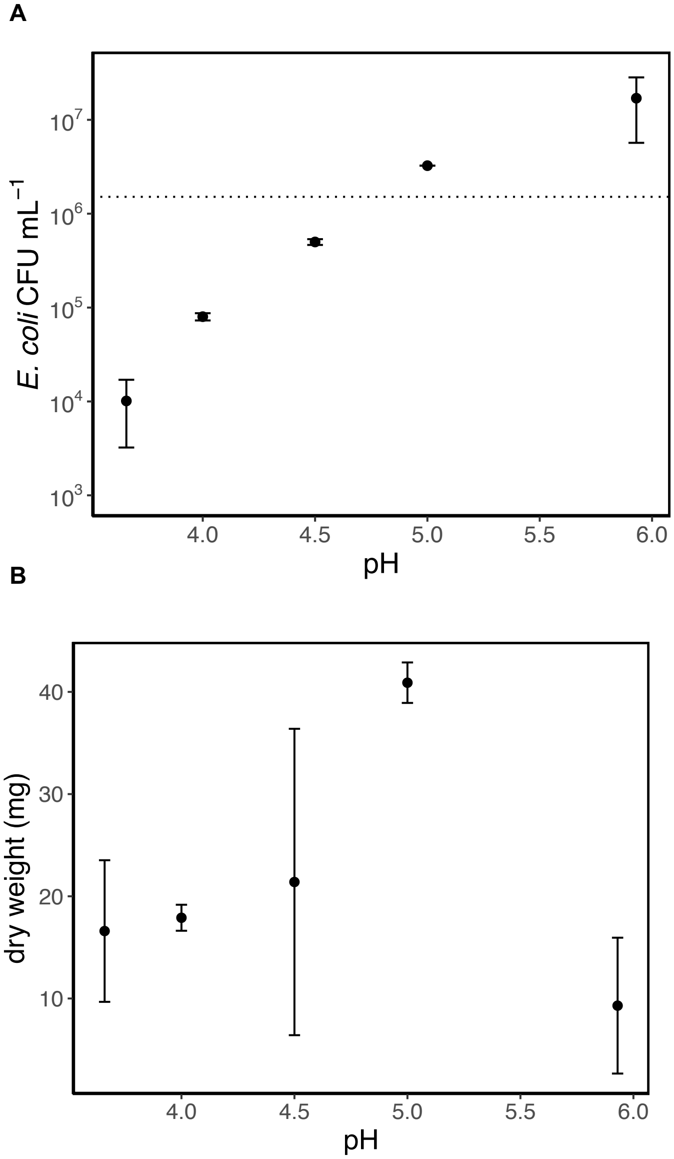


**Fig. S1. Screen to determine optimal pH of HS media for co-culture.** (a) 10mL HS media in a Falcon tube was inoculated with *G. hansenii* and *E. coli*. The pH of the HS media was pre-adjusted to various levels using citric acid (unadjusted HS media had pH of 5.9). After 5 days of incubation, the *E. coli* concentration in the liquid underlying the cellulose pellicle was measured by plating on selective LB-agar. The dotted line indicates the initial concentration of *E. coli.* (b) Dry weight of cellulose pellicles following the 5-day co-culture period.

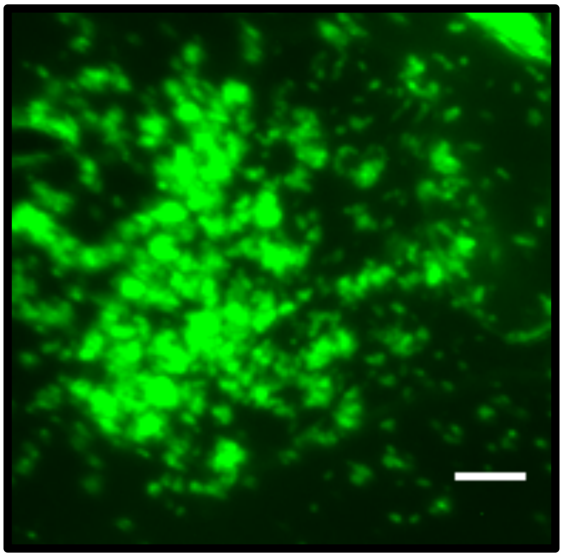

**Fig. S2. Cross-sectional image of capsule containing GFP sequestered by NbGFP-displaying curli fibers.** Section was imaged using an EVOS fluorescent microscope with a GFP light cube (470/22 nm excitation; 525/50 nm emission). Scale bar = 10 μm.

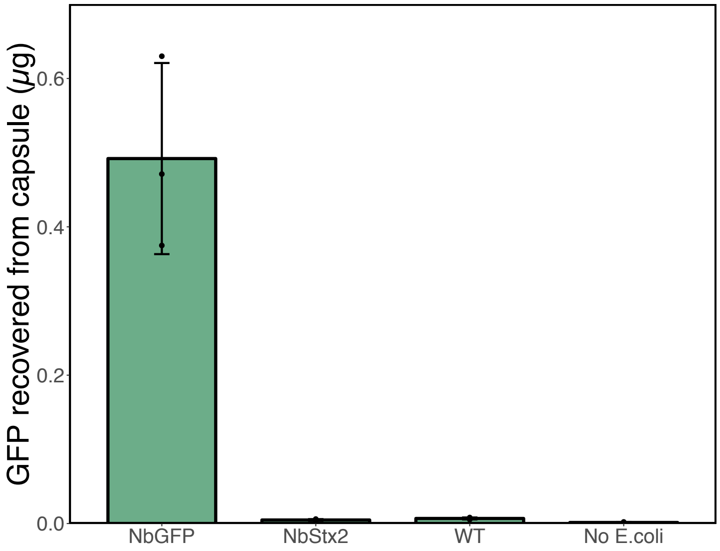


**Fig. S3. Recovery of sequestered GFP from capsules.** Capsules containing sequestered GFP were washed and incubated in 1 M urea with agitation overnight to disrupt the interaction between NbGFP and captured GFP. GFP fluorescence in the 1 M urea supernatant was then measured on a BioTek Synergy NEO plate reader (ex,em = 485,528 nm) to estimate amount of GFP which had been released from the capsules.

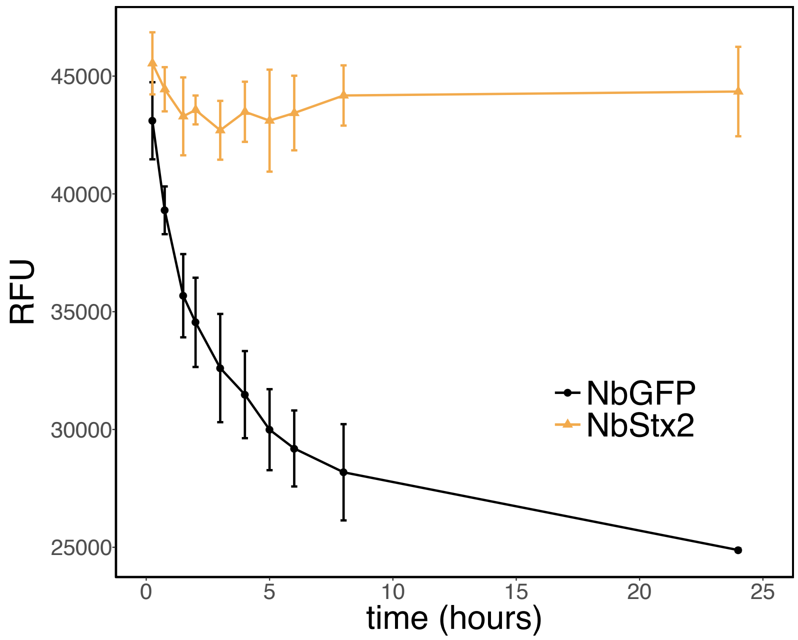


**Fig. S4. Kinetics of GFP sequestration in capsules.** Capsules containing engineered curli fibers with NbGFP fusion domains (or NbStx2 in the negative control) were incubated 0.5 mL PBS containing 4.6 µg/mL purified GFP at 37 ºC with agitation. The supernatant of the solution was sampled with replacement at various time points and GFP fluorescence was measured on BioTek Synergy NEO plate reader (ex,em = 485,528 nm). The fluorescence of the supernatant decreased for the NbGFP sample due to GFP being sequestered in the capsule.

**A**

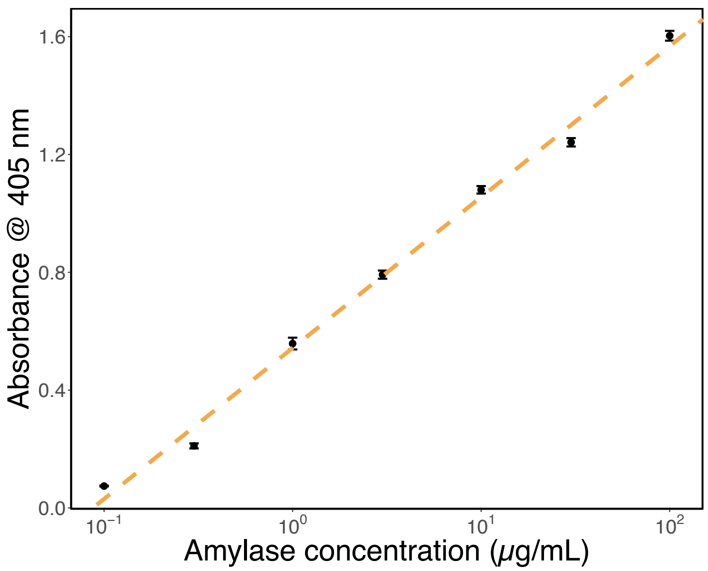

B

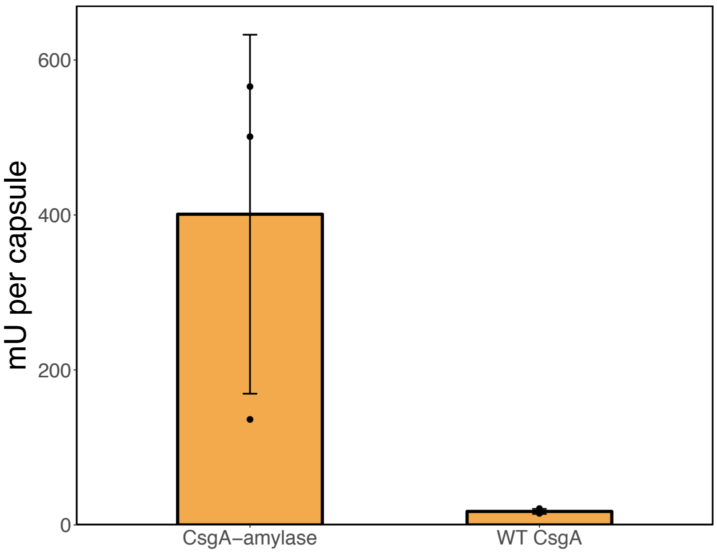


**Fig. S5. Assay to confirm α-amylase activity in capsules containing amylase-displaying curli fibers.** (a) Calibration curve of purified α-amylase computed by measuring absorbance of solution at 405 nm (BioTek Synergy NEO) after co-incubation of amylase and 3 mg/mL chromogenic substrate 4-Nitrophenyl α-D-maltohexaoside under agitation at 37 ºC for 5 hours. (b) α-Amylase activity of capsules containing either amylase-displaying or wild-type curli fibers, computed using calibration curve above.

### Table S1. Strains used in this study.

| Strain | Description | Source |
|---|---|---|
| BL21 | Commercially available protein expression strain | Sigma-Aldrich CMC0016 |
| PQN4 | MC4100, ΔcsgBACEFG, λ(DE3), CamR | Dorval Courchesne *et al*. (*17*) |
| *Gluconacetobacter hansenii* | Acetic acid bacterium which produces high levels of BC | ATCC 53582 |
| HB101 | Commercially available protein expression strain | Promega L2015 |

### Table S2. Plasmids used in this study.

| Plasmid | Description | Source |
| --- | --- | --- |
| pBbA8k-RFP | Arabinose-inducible expression of mRFP1 | Addgene # 35273 |
| pBbB8k-csg-WT | Arabinose-inducible expression of the full *csg* operon. *Csg* operon inserted in place of GFP in Addgene # 35363**.** | This study |
| pBbB8k-csg-NbGFP | Arabinose-inducible expression of the full *csg* operon in which *csgA* is fused to NbGFP, a VHH domain specific for GFP, by a 14aa GS linker. | This study |
| pBbB8k-csg-NbStx2 | Arabinose-inducible expression of the full *csg* operon in which *csgA* is fused to NbStx2, a VHH domain specific for shiga toxin 2, by a 14aa GS linker. | This study |
| pBbB8k-csg-amylase | Arabinose-inducible expression of the full *csg* operon in which is *csgA* fused to α-amylase from *Bacillus licheniformis* by a 14aa GS linker. | This study |
| pLO7 | Constitutive expression of *luxCDABEFG* operon from *V. harveyii*. | This study |
| pBR322-Ure | Urease gene cluster from *S. pasteurii*. | Liang et al. (*49*) |
